# Rapid rewiring of an archaeal transcription factor function via flexible cis-trans interactions

**DOI:** 10.1101/2024.11.20.624553

**Authors:** Mar Martinez Pastor, Cynthia L. Darnell, Angie Vreugdenhil, Amy K. Schmid

## Abstract

For microbial cells, an appropriate response to changing environmental conditions is critical for viability. Transcription regulatory proteins, or transcription factors (TFs) sense environmental signals to change gene expression. However, it remains unclear how TFs and their corresponding gene regulatory networks are selected over evolutionary time scales. The function of TFs and how they evolve are particularly understudied in archaeal organisms. Here we identified, characterized, and compared the function of the RosR transcription factor across three related hypersaline adapted archaeal model species. RosR was previously characterized as a global regulator of gene expression during oxidative stress in the species *Halobacterium salinarum* (*hs*RosR). Here we use functional genomics and quantitative phenotyping to demonstrate that, despite strong sequence conservation of RosR across species, its function diverges substantially. Surprisingly, RosR in *Haloferax volcanii* (*hv*RosR) and *Haloferax mediterranei* (*hm*RosR) regulates genes whose products function in motility and outer membrane structure, leading to significant defects in motility when *rosR* is deleted. Given weak conservation and degeneration in cis-regulatory sequences recognized by the RosR TF across species, we hypothesize that the RosR regulatory network is readily rewired during evolution across related species of archaea.

**SIGNIFICANCE STATEMENT:**

- Gene regulation enables cells to sense and respond to environmental signals. The mechanisms by which gene regulatory circuits change and adapt over evolutionary time scales remain unclear, especially in understudied domains of life like the archaea.
- Here we demonstrate that the archaeal-specific RosR transcription protein plays fundamentally different roles in environmental response of related archaeal species (oxidative stress protection vs motility). This divergence likely occurred through reduced selectivity of RosR for certain DNA sequence motifs.
- These findings are surprising given strong sequence and structural conservation of RosR and suggest that transcription regulator function can diverge rapidly through flexible protein-DNA interactions. This mechanism of divergence is shared between eukaryotes and archaea, suggesting ancient origins.

Subject categories: Bioinformatics, microbiology, genetics, gene regulatory networks

## INTRODUCTION

Transcription regulatory networks (TRNs) across life have been observed to “rewire” across evolutionary time scales, whereby transcription factor proteins (TFs) regulate different gene targets and therefore affect cellular functions (Perez and Groisman, 2009b, a; Hackley *et al*., 2024b; Martinez Pastor *et al*., 2024). Cells use TRNs to change gene expression in response to environmental signals. TFs bind to promoters to recruit or inhibit RNA polymerase, thereby activating or repressing transcription, respectively. TRNs comprised of many TFs (trans factors) regulate their target genes by binding to regulatory sequences proximal to gene promoters (cis factors). TRNs are crucial across life for essential cellular processes ranging from stress response to cell development (Martinez-Pastor *et al*., 2017b). Theoretically, TRN rewiring can occur through mutations in cis or trans factors. In eukaryotes, rewiring is thought to occur primarily through cis-regulatory mutations across related species (Johnson, 2017), although significant trans rewiring of TF functions has also been observed (Dalal *et al*., 2016). In contrast, trans factors appear to be a primary mode of TRN rewiring in bacteria through horizontal gain and loss of TF genes (Perez and Groisman, 2009b, a). However, whether such rewiring is adaptive needs additional clarity, and it is not yet known whether general mechanistic rules govern network rewiring across all domains of life. Archaea are a particularly understudied domain of life, especially from the standpoint of TRN architecture, function, and rewiring across evolutionary time.

Although understudied, archaea play important roles in the human microbiome, global nutrient and energy cycles, and biotechnological applications (Jarrell *et al*., 2011). Importantly, archaeal protein families are thought to be ancestral to all domains of life (Kim and Caetano-Anolles, 2012), but this is coupled with substantial horizontal gene transfer between archaea and bacteria (Garcia-Vallve *et al*., 2000). The archaeal transcription information processing machinery exemplifies these interwoven evolutionary processes (Kyrpides and Ouzounis, 1999; Perez-Rueda and Janga, 2010; Fouqueau *et al*., 2018). Archaeal transcription proteins that recruit RNA polymerase to promoters are ancestral and homologous to those of eukaryotes, especially TATA-binding protein (TBP) and transcription factor B (TFB) that assemble the pre-initiation complex on DNA to set the basal level of transcription (Bell *et al*., 2001). In contrast, regulation of transcription in response to environmental cues depends on TFs homologous to (and perhaps transferred horizontally from) those of bacteria (Perez-Rueda and Janga, 2010; Lemmens *et al*., 2019). These TFs typically possess a helix-turn-helix DNA binding domain linked to a ligand-binding activation domain (Perez-Rueda and Janga, 2010), although a few single-domain DNA binding TFs have recently been characterized in archaea (Kutnowski *et al*., 2018; Darnell *et al*., 2020; Liao *et al*., 2021; Mondragon *et al*., 2022).

Many archaeal model species for molecular and cellular biology are well known as stress response specialists, dominating the biomass in extreme environments from boiling hot springs to saturated salt lakes (Schmid *et al*., 2020). Salt adapted haloarchaea reside in salt flats and lakes that can reach sodium chloride saturation, where they experience high levels of oxidative stress (Andrei *et al*., 2012; Jones and Baxter, 2017). They provide a unique suite of experimentally tractable archaeal model organisms for comparative evolutionary studies of transcription mechanisms and TRNs that respond to stress (Allers *et al*., 2010; Leigh *et al*., 2011; Martinez-Pastor *et al*., 2017b). In particular, the organisms of interest here (*Haloferax volcanii, Haloferax mediterranei* and *Halobacterium salinarum*) vary in the optimum salinity level for growth (∼2.5 M – 3.5 M NaCl, respectively) but encode many homologous TFs (Ng *et al*., 2000; Hartman *et al*., 2010; Han *et al*., 2012). Recent comparative TRN research suggests that rewiring across related species of haloarchaea can occur via acquisition of novel TFs [i.e. rewired trans factors, (Martinez-Pastor *et al*., 2017a; Hackley *et al*., 2024a; Martinez Pastor *et al*., 2024)]. Although the cis-regulatory consensus sequences for each TF are strongly conserved in general, TF-target gene connections appear to change through wholesale deletion or acquisition of TFs and/or cis sequences (Hackley *et al*., 2024a; Hackley *et al*., 2024b). However, it remains unclear in archaea whether these recently published examples are general to all archaeal TRN rewiring mechanisms.

The PadR family archaeal-specific transcription factor RosR, first characterized as an oxidative stress regulator in *Halobacterium salinarum* (Sharma et al., 2012; Tonner et al., 2015), serves as a paradigmatic case study for TRNs in archaea. *hs*RosR (VNG_0258H, NCBI ID VNG_RS01050) regulates the transcription of over 300 genes in *Hbt. salinarum*, facilitating the organism’s resistance to oxidative stressors such as hydrogen peroxide and paraquat (Sharma *et al*., 2012). *hs*RosR primarily represses its gene targets during standard conditions, then releases DNA to de-repress expression in multiple waves (early, middle, late gene clusters) over the course of 60 minutes upon oxidant exposure (Tonner *et al*., 2015). The mechanism of *hs*RosR-DNA dissociation remains unknown. Gene targets include catalase, peroxidase, and over 20 TF-coding genes, including TFBs. These TFs feed back to regulate the expression of the gene encoding *hsrosR* and each other (Tonner *et al*., 2015). This multifaceted regulatory role of RosR highlights the intricate interplay between TFs and their target genes in archaeal adaptation to extreme environments.

However, despite the extensive characterization of *hs*RosR in *Hbt. salinarum*, the regulatory roles of its homologous proteins in other halophilic species remain poorly understood. Notably, *Haloferax volcanii* and *Haloferax mediterranei* are both halophiles adapted to more moderately saline environments (∼2.5M), and both encode RosR homologs (HVO_0730 and HFX_0688, respectively) of *hs*RosR (Sharma *et al*., 2012). Curiously, *hv*RosR is not induced in response to peroxide or hypochlorite stress in *Hfx. volcanii*, suggesting that RosR may play different roles in related species of haloarchaea (Gelsinger and DiRuggiero, 2018; McMillan *et al*., 2018). These preliminary findings underscore the need for comparative analysis of RosR across halophilic species, and position RosR as an excellent model system for investigating network rewiring across species.

Here we used genomics, genetics, and quantitative phenotyping to show that, despite strong structural, sequence, and binding motif conservation of RosR across haloarchaea, the genes it controls diverge strongly between species living in different salinity regimes. Surprisingly, RosR is not required for growth under oxidative stress in either *Hfx*. species. Rather, *hm*RosR and *hv*RosR each regulate a small number of genes encoding membrane functions and archaellum machinery, respectively. This function stands in sharp contrast to the global regulatory function of *hs*RosR in *Hbt. salinarum* oxidative stress response. Taken together, these results suggest that the RosR transcriptional landscape is highly plastic during evolution.

## RESULTS

### RosR is a PadR family homolog that is strongly conserved throughout Haloarchaea

Previously we demonstrated that RosR is a PadR family (PFAM 03551) winged helix-turn-helix (wHTH) TF conserved at the amino acid level across selected haloarchaea, including *Hfx. volcanii* (Sharma et al., 2012). Given the recent expansion in bacterial and archaeal genome sequencing, we extended this sequence analysis to determine the phylogenetic conservation across the PadR family. Protein sequences from 767 bacterial, viral, and haloarchaeal genomes were drawn from the PFAM database (Table S1). Unrooted neighbor joining tree topology showed that haloarchaeal RosR sequences formed a distinct monophyletic and recently radiating clade, suggesting strong RosR TF family sequence conservation across haloarchaea (Figure 1A).

**Figure 1.**
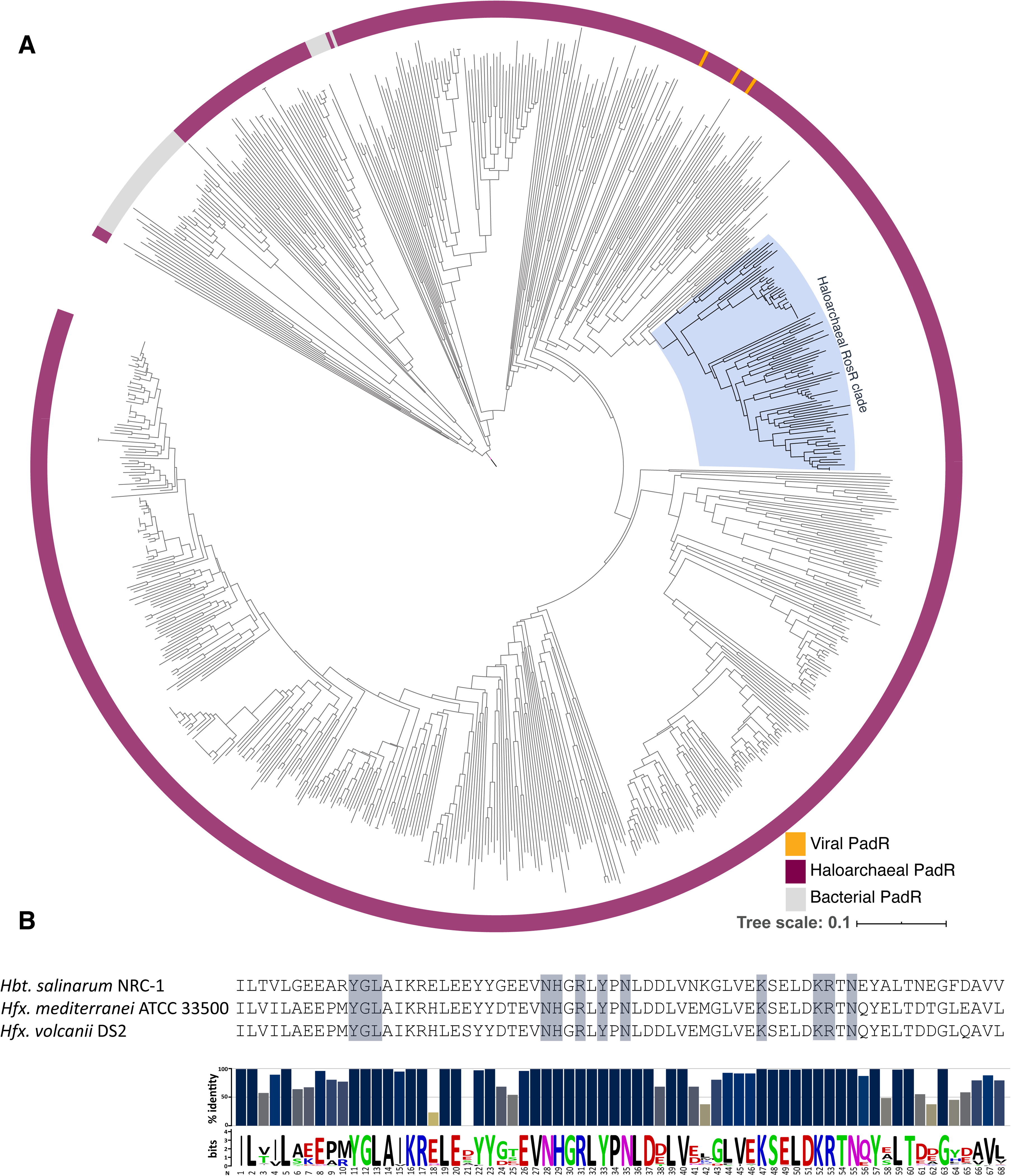
RosR is highly conserved throughout Haloarchaea. (A) Haloarchaeal clade within larger PadR tree (See also Supplemental Figure 1 for a list of sequences included). Magenta outer ring denotes haloarchaeal proteins; gray denotes bacterial proteins; orange denotes archaeal virus protein. Blue shaded region indicates RosR homologs by amino acid sequence and genomic synteny. (B) Alignment of the wing and alpha helical DNA binding regions of the RosR homologs from the three species of interest (top panel). Highlighted regions indicate residues that contact DNA in *hs*RosR *in vitro* [in the wing (R74, corresponds to residue 53 in the weblogo at bottom) and the helix (H50, corresponds to residue 29), (Kutnowski *et al*., 2018; Kutnowski *et al*., 2019)]. The haloarchaeal consensus sequence for the RosR DNA binding domain (wing and helix) was determined through aligning the 90 RosR homologs and calculating percent identity (middle panel) and generating a weblogo (bottom panel).

We next investigated RosR primary sequence homology in greater detail across the haloarchaeal model species of interest here. The amino acid sequence of RosR from *Halobacterium salinarum* (*hs*RosR, VNG_0258H) was used to query the genomes for each of *Haloferax volcanii* (Hartman et al., 2010) and *Haloferax mediterranei* (Han et al., 2012), followed by reciprocal BLAST. This identified HVO_0730 in *Hfx. volcanii* (NCBI gene identifier HVO_RS08205) which showed 65.22% sequence identity to *hs*RosR and 91.6% to *Hfx. mediterranei,* HFX_0688 (HFX_RS03340, *hm*RosR). *hm*RosR also exhibited 65.22% identity to *hs*RosR (see also Table S2 for additional database identifier details). Both HVO_0730 and HFX_0688 were the top hits to *hs*RosR in each BLAST search and were used in subsequent experiments and analyses. The per-residue identity throughout the helix and wing regions of the protein was extremely high across the three species. Residues previously implicated in DNA binding from the *hs*RosR X-ray crystal structure were conserved (Figure 1B, top (Kutnowski *et al*., 2019)). In addition, 100% conservation was observed at 38 of 68 residues in the DNA binding region (Figure 1B, bottom). Taken together, these results suggest that RosR amino acid sequence is strongly conserved across the haloarchaeal species of interest, especially in the DNA binding region.

### RosR is dispensable under oxidative stress in *Haloferax* species

Because of the strong conservation of RosR, we hypothesized that this TF may be important for oxidative stress survival in *Hfx*. species as was previously shown for *Hbt. salinarum* (Sharma et al., 2012). To test this, we constructed Δ*rosR* in-frame deletion strains in *Hfx. volcanii* (*hv*Δ*rosR*) and *Hfx. mediterranei* (*hm*Δ*rosR*) and confirmed by whole genome resequencing that all copies of the gene had been deleted from the respective genomes (see Methods and Table S2). No second site mutations were observed.

We first determined the amount of oxidant to use for testing Δ*rosR* phenotypes across species. We conducted growth curve experiments in rich medium across titrations of varying concentrations of peroxide (H_2_O_2_) and paraquat (PQ) in the parental strain of each species (Figure 2, Table S3). Each species grew as expected in rich medium without oxidant (standard conditions, *Hfx. volcanii* ∼3 h doubling time; *Hfx. mediterrenei* ∼2 h, Figure 2, Figure S1A). The growth of each *Hfx.* species showed similar sensitivity when peroxide (H_2_O_2_) was added at culture inoculation, with no growth observed for either strain at 2 mM H_2_O_2_ (Figure 2A and B). However, the *Hfx. volcanii* parental strain appeared slightly more susceptible to 0.5 mM H_2_O_2_ (Figure 2A) than *Hfx. mediterranei* (Figure 2B). The *Hfx.* species are more sensitive to peroxide than *Hbt. salinarum*, which exhibits only slight growth inhibition at 5 mM H_2_O_2_ added at inoculation (Sharma *et al*., 2012). Surprisingly, *Hfx mediterranei* was more resistant to paraquat (PQ) treatment than the other species (Figure 2C and D), with some residual growth observed at concentrations as high at 4 mM PQ (Figure 2D). *Hfx. volcanii* was more sensitive, exhibiting growth impairment at 0.67 mM PQ (Figure 2C). *Hbt. salinarum* was the most sensitive of the species to PQ: we previously observed growth inhibition with as little as 0.33 mM PQ (Sharma *et al*., 2012). Taken together, these data suggest that both *Hfx.* species are more susceptible to peroxide stress than *Hbt. salinarum* but less sensitive to PQ. The concentrations causing slight impairment of growth in each parent strain were chosen to test for Δ*rosR* growth phenotypes (concentrations are bounded by a box in Figure 2).

**Figure 2.**
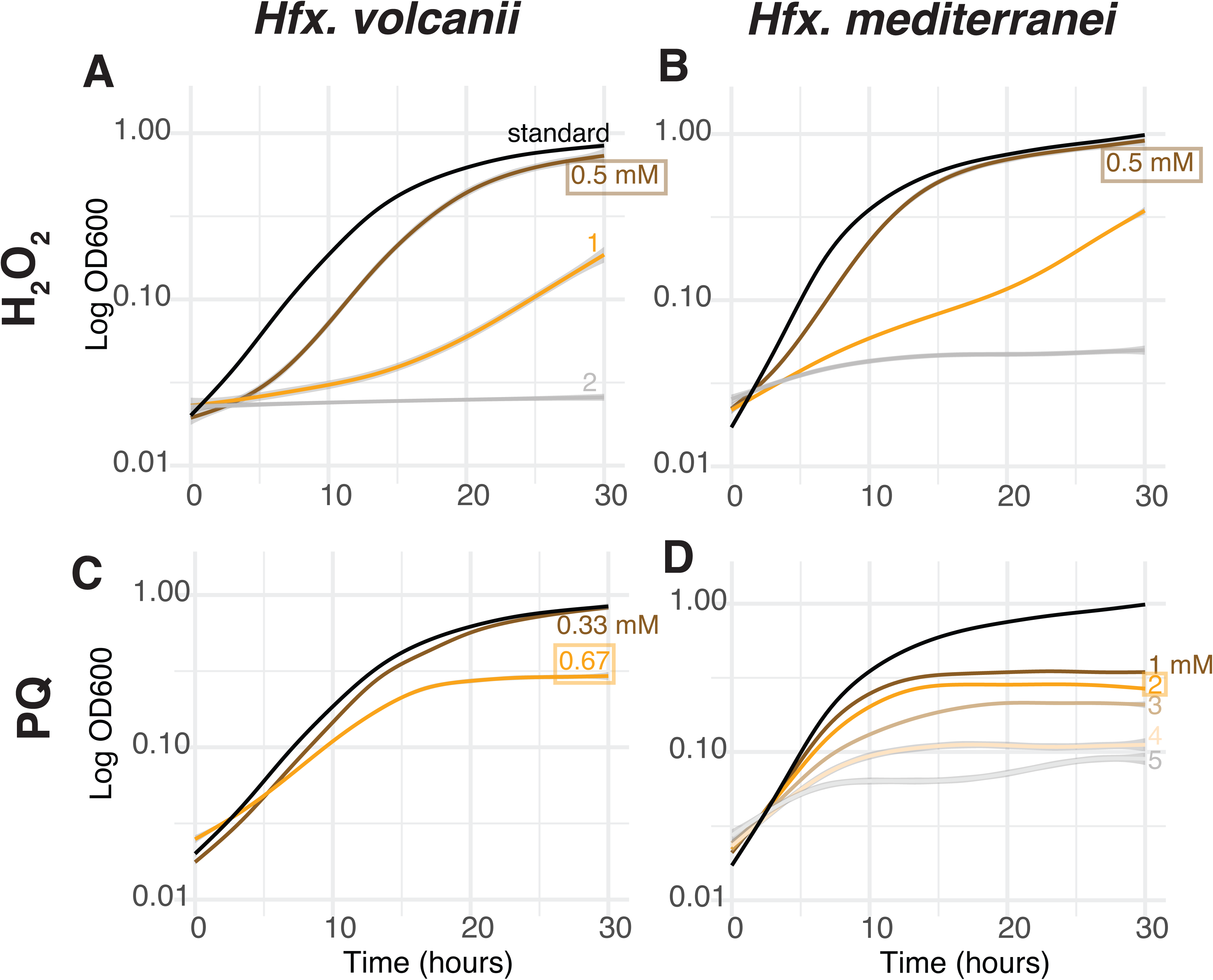
Haloarchaeal species exhibit differential susceptibility to oxidative stress conditions. Growth is plotted as the log10 optical density at 600 nm (OD600) over time (hours). Each curve represents the generalized additive model (GAM) fit to the raw data from at least 3 biological replicate experiments (seeded from starter cultures inoculated with separate colonies), each with 3 technical replicates (aliquots from the same starter culture). Shaded grey ribbons indicate the standard error. Error is low where ribbons are not visible. Concentrations are written above the corresponding growth curve in each panel. Concentrations surrounded by a box is that condition chosen for testing Δ*rosR* growth in subsequent experiments. (A) *Hfx. volcanii* growth under a titration of peroxide (H_2_O_2_) concentrations. (B) *Hfx. mediterranei* growth under H_2_O_2_. (C) *Hfx. volcanii* growth under varying paraquat (PQ) concentrations. (D) *Hfx. mediterranei* growth under PQ.

Using these concentrations of oxidant, we conducted growth assays for *hm*Δ*rosR* and *hv*Δ*rosR* compared to the known growth defect of *hs*Δ*rosR* (Sharma 2012). We recapitulated the strong growth impairment of *hs*Δ*rosR* under paraquat and 5 mM H_2_O_2_ conditions relative to the *Hbt. salinarum* parent control strain (Figure 3, left; Figure S1B). In contrast, no growth impairment was observed for *hm*Δ*rosR* in at 0.5 mM H_2_O_2_ (Figure 3, middle and right; Figure S1B; Table S3). *hv*Δ*rosR* showed a slight growth impairment under 0.5 mM H_2_O_2_, but this was found not to be statistically significant according to the area under the curve metric (AUC, (Todor *et al*., 2014), Welch’s two-sample t-test *p* = 0.1869, Figure S1C). Similarly, no growth defect was observed for Δ*rosR* in either *Hfx.* species under paraquat conditions (Figure 3, lower panels, Figure S1B and C). Taken together, these data surprisingly suggest that, despite strong sequence homology across species, RosR plays a cellular role other than oxidative stress response in the *Hfx.* species.

**Figure 3.**
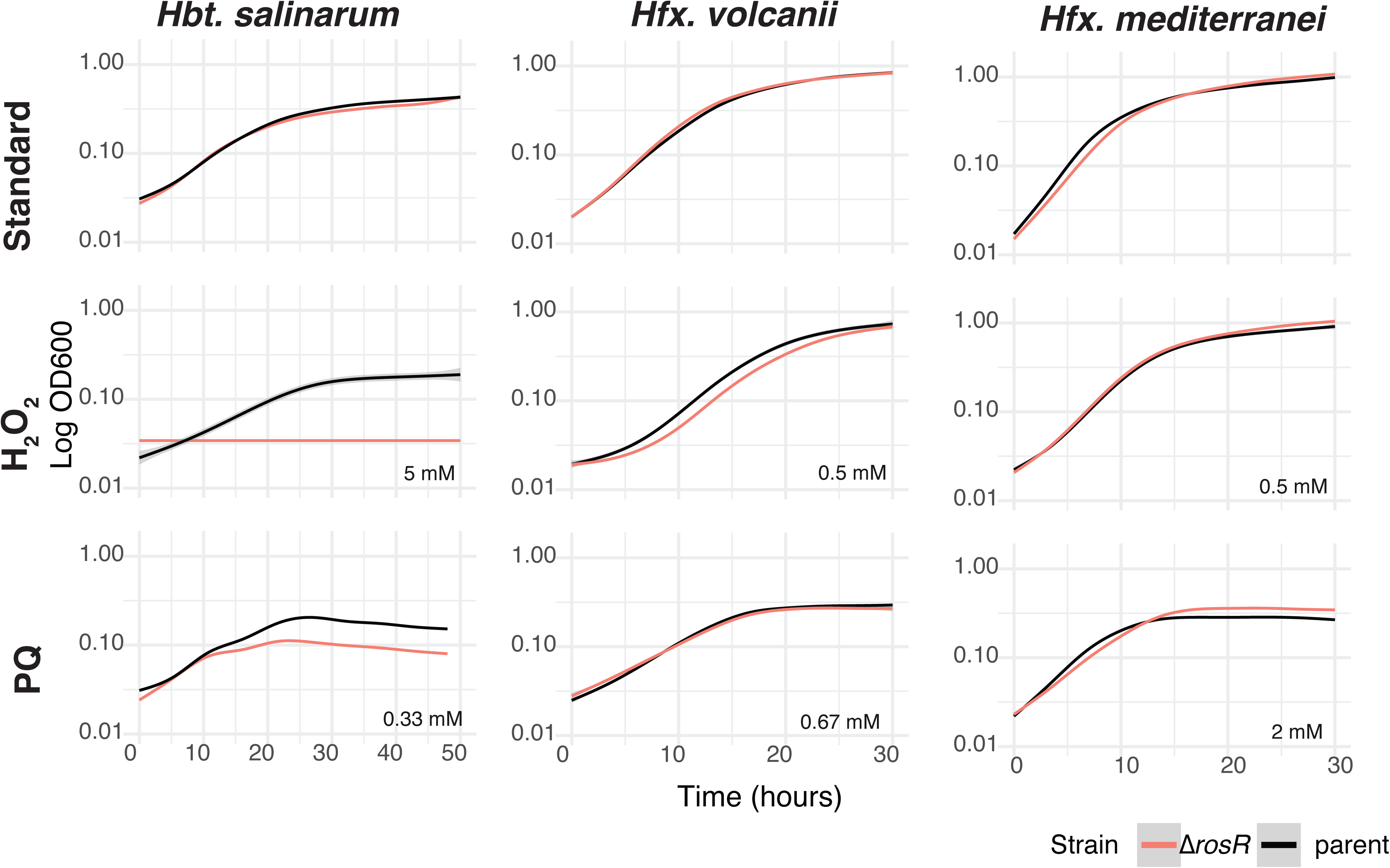
RosR is dispensable for growth under oxidative stress conditions in *Hfx*. species. Growth is plotted as the log10 optical density at 600 nm (OD600) over time (hours). As in Figure 2, each curve represents the generalized additive model (GAM) fit to the raw data from at least 3 biological replicate experiments, each with 3 technical replicates. Shaded grey ribbons indicate the standard error. Error is low where ribbons are not visible. Concentrations are written at the lower right corner of each growth curve panel. Pink ribbons represent Δ*rosR* growth, black represent parent strain (see also legend at lower right). Left column, recapitulation of previously reported growth phenotypes for *hs*Δ*rosR* in *Hbt. salinarum* (for the purpose of comparison (Sharma *et al*., 2012)). Middle column, *hv*Δ*rosR* growth compared with the parent strain; right column, *hm*Δ*rosR* growth compared with the parent strain.

### In *Hfx* species, RosR regulates the expression of cell surface and motility genes

To determine possible functions for RosR in *Hfx. mediterranei* and *Hfx. volcanii* we conducted functional genomics. Transcriptomes were measured by RNA sequencing transcriptomics (RNA-seq) in Δ*rosR* vs parent strains. Genome-wide binding site location analysis was conducted using chromatin immunoprecipitation coupled to sequencing (ChIP-seq). Both experiments were conducted in standard growth conditions (mid-exponential phase in rich medium) to enable comparisons with known binding sites for *hs*RosR, which binds and represses H_2_O_2_-responsive genes in standard conditions (Tonner *et al*., 2015).

In *Hfx. volcanii*, 74 sites genome-wide were enriched for RosR binding in ChIP-seq data, and 16 genes were significantly differentially expressed in *hv*Δ*rosR* relative to the parent strain in RNA-seq (6 genes downregulated and 10 upregulated, Table S4). Four loci were detected in the intersection of these two datasets, but only *arlA2* (HVO_1211, HVO_RS10520) passed further visual inspection of the ChIP-seq data (Table S4). *arlA2* encodes an archaellin protein, a known structural component of the archaeal motility apparatus (Figure 4A (Tripepi *et al*., 2010)). RosR bound to a region overlapping the 3’ end of *arlA2* and the intergenic space downstream of the convergently transcribed *cirA* (Figure 4A, Table S4), a gene recently shown to encode a regulator of motility (Chatterjee *et al*., 2024). Both *arlA2* and *cirA* are reside within the large gene cluster encoding motility functions such as the archaellum and chemosensory (*che*) proteins (Figure 4A, top). *arlA1* and *arlA2* were significantly downregulated in *hv*Δ*rosR* relative to the parent strain, but otherwise *hvrosR* deletion had modest effects on the transcriptome (Figure 4B). However, it remains unclear how *Hfx. volcanii* RosR activates the expression of *arlA2* by binding the 3’ end of the gene. Because ChIP sequencing read depth was low in this region relative to what is expected for ChIP-seq binding enrichment peaks in this organism (Figure 4A, (Hackley *et al*., 2024a; Martinez Pastor *et al*., 2024)), we validated by ChIP coupled to quantitative PCR (ChIP-qPCR) using three different amplicons (Figure 4C). ChIP-qPCR showed that RosR binding was statistically significantly enriched relative to the parent strain at each of three ∼100 bp regions within this locus (Figure 4C). A putative *hv*RosR cis-regulatory binding sequence motif similar to the previously reported TGACA half-site of *hs*RosR (Kutnowski *et al*., 2019) was discovered using MEME with input from *hv*RosR ChIP-seq enriched sequences (Figure 4D, Table S4, (Bailey *et al*., 2015)). A similar motif was detected in 20 locations genome-wide (Table S4). 14 of these motifs were located within 250 bp of the start of *hv*RosR ChIP-seq peaks, including three genes encoding other TFs (e.g. TFB HVO_1478). However, genes nearby these peaks were not differentially expressed in the *hv*Δ*rosR* strain relative to the parent (Table S4). No cis-regulatory sequence motifs were detected near the *arlA2* binding site. Together these data suggest that: (a) the *hv*RosR binding site is quite degenerate and difficult to discern by computational methods alone; and/or (b) *hv*RosR may bind genes without regulating their expression (i.e. may require another factor to change gene expression at binding sites). Nevertheless, these results support the hypothesis that *hv*RosR is necessary for activation of the *arlA1* and *arlA2* genes, whose products function in motility in *Hfx. volcanii*.

**Figure 4.**
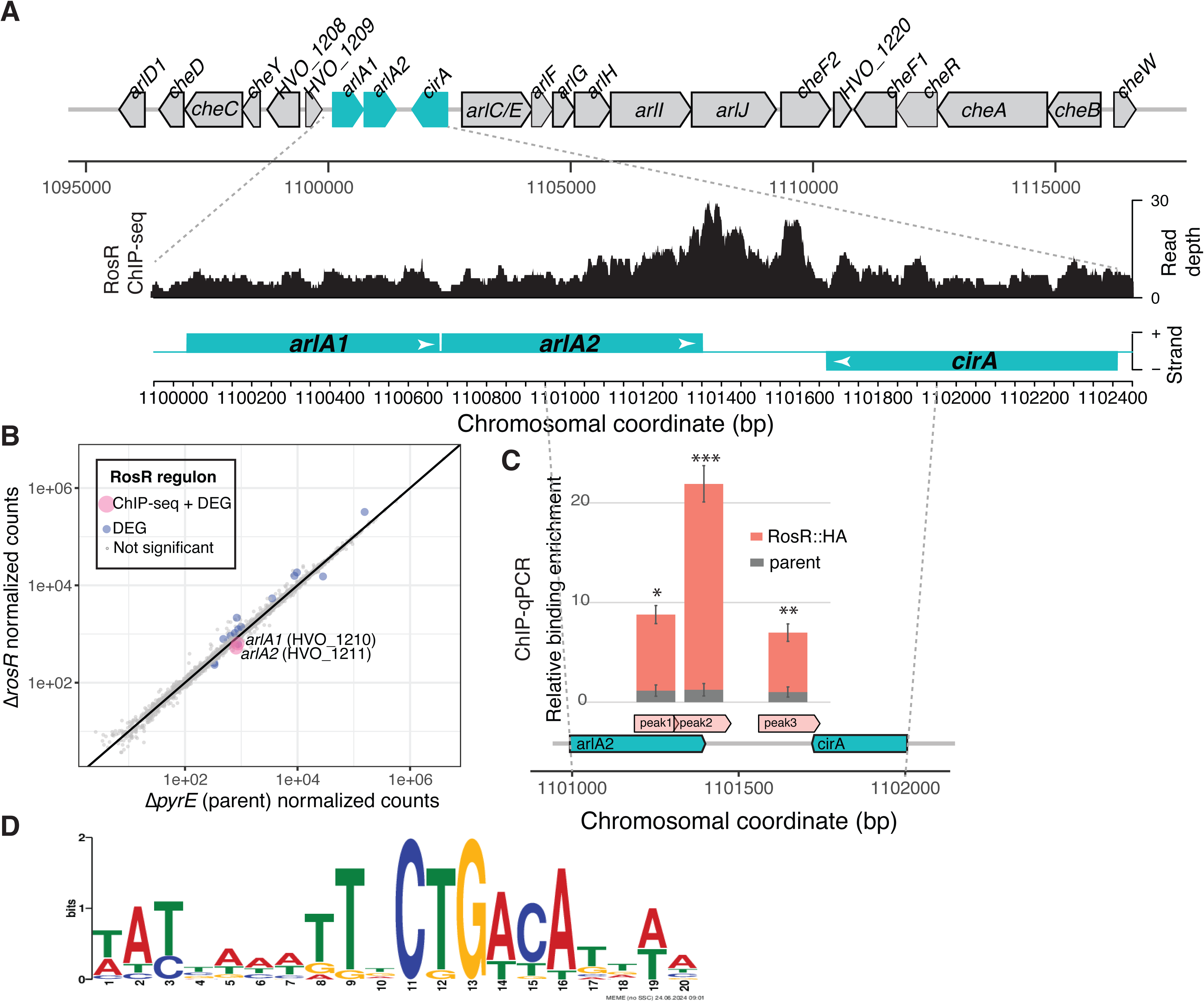
*hv*RosR activates *arlA1* and *arlA2*, encoding structural components of the archaellum in *Hfx. volcanii*. (A) Top, genomic region encoding the archaellum and related motility functions; middle, ChIP-seq data for the region highlighted in teal (read depth y-axis is shown at right); bottom, chromosomal coordinates for genes in the zoomed region, with genes on the forward strand depicted on top of the line and reverse strand below the line. (B) Scatterplot of normalized counts from RNA-seq data, with parental control strain counts on the X-axis and *hv*Δ*rosR* counts on the y-axis. Genes passing the significance threshold of p < 0.05 and log2 fold change >= |1| are shown in blue, those significant genes also bound in ChIP-seq data in pink (see legend). (C) ChIP-qPCR validation of ChIP-seq data. Bar height represents the mean enrichment of *hv*RosR binding each site relative to a control region. Error bars represent the standard deviation from the mean of triplicate samples. *hv*RosR-HA enrichment is shown in salmon and PyrE2-HA parent strain control in grey. Amplicons tested are shown in orange below the bar graph (“peak 1, peak 2, peak 3”) and are set relative to chromosomal position of the corresponding genes. Asterisks indicate the statistical significance of enrichment for each peak (salmon bars) relative to the parent control (grey bars) by two-sided unpaired Student’s t-test, \**p* < 1.02 x 10^-3^, ** *p* < 4.71 x 10^-4^, \*\*\**p* < 6.31 x 10^-5^. (D) Logo of the consensus binding motif detected computationally from *hv*RosR ChIP-seq binding site sequences. Motif position in nucleotides is given on the x-axis and bit score of per-base representation in the position weight matrix is given on the y-axis. See also Supplementary Table S4 for detailed RNA-seq, ChIP-seq, and motif data.

In *Hfx. mediterranei*, *hm*RosR binding enrichment was detected at 32 genomic sites by ChIP-seq and 150 genes were significantly differentially expressed in *hm*Δ*rosR* relative to the parent strain (91 downregulated and 59 upregulated, Table S4). Of the genes detected as significant in both ChIP-seq and RNA-seq datasets, two genes, HFX_0465 and HFX_1425, also had detectable cis-regulatory motifs upstream (Figure 5A and B, respectively). This TGACA-N6-TGACA direct repeat motif was detected computationally using MEME (Figure 5C, Table S4, Methods). Three other genes were detected as significant in both the ChIP-seq and RNA-seq datasets (HFX_1711, HFX_1971, HFX_1666), but no cis-regulatory binding motif was detected nearby (Figure 5D). The cis-sequence motif was detected within each of the ChIP-seq peaks upstream of HFX_0465 and HFX_1425 (Figure 5A and B, red ticks), and was also detected 16 other times throughout the *Hfx.* mediterranei genome (Table S4). RNA-seq transcriptomics demonstrated that HFX_1425 is activated by *hm*RosR under standard conditions, whereas HFX_0465 is repressed (Figure 5D). The HFX_0465 gene neighborhood is predicted to encode cold shock proteins and a signal peptidase (Figure 4A, right). HFX_0465 encodes a predicted sulfatase that exhibits 46% amino acid sequence identity to *Hfx. volcanii* HVO_0069, mutants of which exhibit motility defects (Legerme *et al*., 2016). HFX_1425 encodes a predicted PGF domain protein with 32% identity to HVO_2160, which encodes membrane protein rod-determining factor A, RdfA (Schiller *et al*., 2024). Taken together, these data suggest that *hm*RosR plays a role in cell surface and/or membrane function and possibly motility.

**Figure 5.**
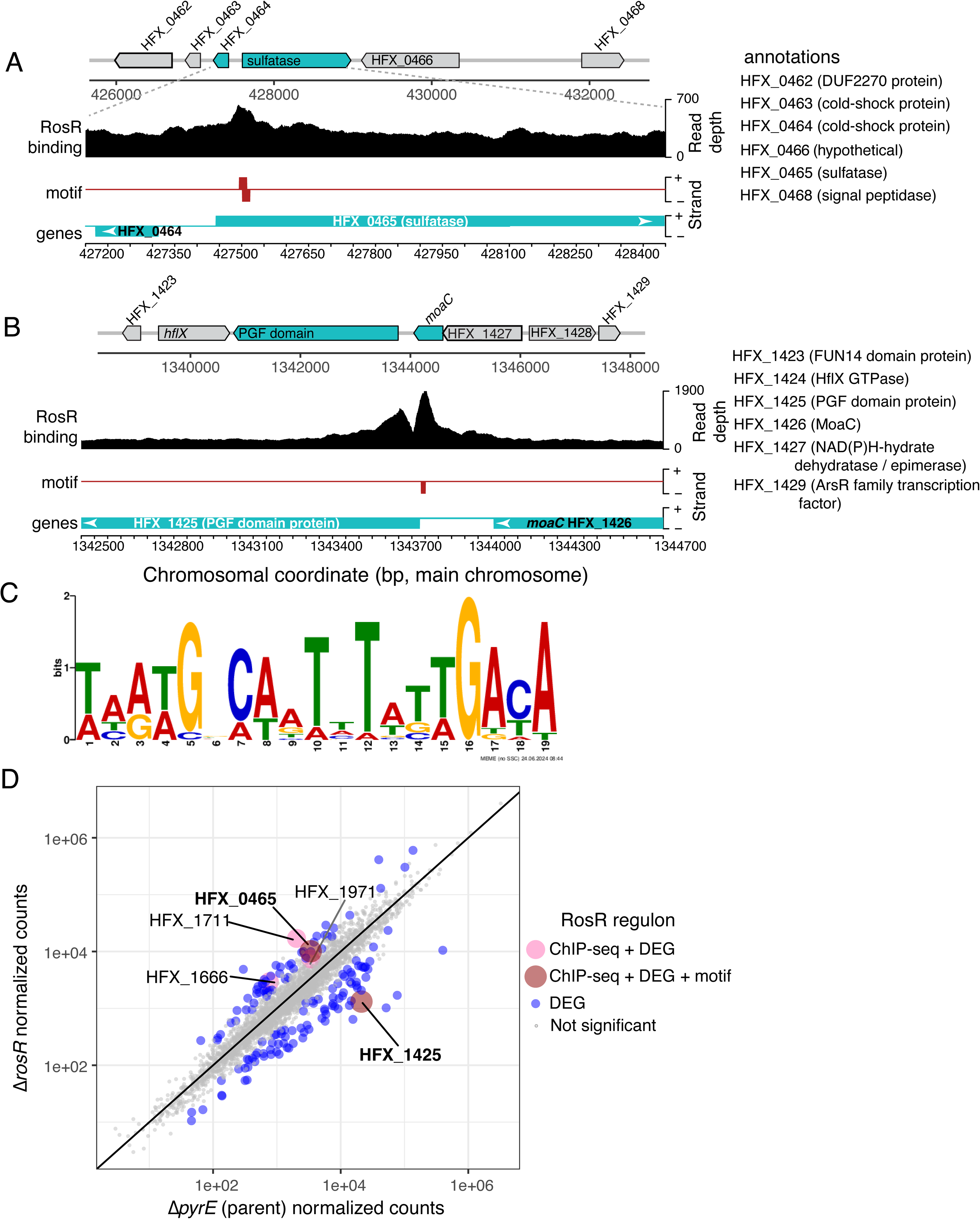
*hm*RosR regulates genes with membrane-related functions in *Hfx. mediterranei*. (A) ChIP-seq binding data, cis-regulatory motif position, and genomic context for HFX_0465 (sulfatase). Top: zoomed-out genomic region with genes of interest in teal. Chromosomal coordinate in bp is given on the x-axis. Second track from the top shows ChIP-seq read depth in black, third track shows position of cis-regulatory motif in red, bottom track shows the genes again zoomed in. Annotations for all genes in the region of interest are listed at the far right. (B) ChIP-seq binding data, cis-regulatory motif position, and genomic context for HFX_1425 (encoding PGF-domain containing protein). Colors and tracks are designated as in panel A. (C) Logo of the consensus binding motif detected computationally from RosR ChIP-seq binding site sequences. Motif position in nucleotides is given on the x-axis and bit score of per-base representation in the position weight matrix is given on the y-axis. (D) Scatterplot of normalized counts from RNA-seq data, with parental control strain counts on the X-axis and Δ*rosR* counts on the y-axis. Genes passing the significance threshold of p < 0.05 and log2 fold change >= |1| are shown in blue, those significant genes also bound in ChIP-seq data in pink, significant genes bound in ChIP-seq that also contain a predicted motif in brick red (see legend). Supplementary Table S4 lists detailed data for RNA-seq, ChIP-seq, and motif data.

### *hm*RosR and *hv*RosR are required for cell motility

Based on the functional genomics data (Figure 4 and 5), we hypothesized that RosR may play a role in motility in the *Hfx.* species. We therefore conducted motility assays in which each of the *hv*Δ*rosR* and *hm*Δr*osR* strains were stabbed into soft agar and the swimming halo area was measured (Methods). As expected from previous functional genomics studies in *Hbt. salinarum* in which RosR does not regulate motility genes, *hs*Δ*rosR* motility halo area was indistinguishable from that of the Δ*ura3* parent strain (p = 0.259, Figure 6A and Figure S2). *hs*Δ*rosR* was also significantly more motile than the negative control strain Δ*aglB*, which was previously shown to be non-motile (*p* < 4.68 x 10^-5^)(Zaretsky *et al*., 2019). In contrast, *hv*Δ*rosR* and *hm*Δ*rosR* motility halos were both significantly smaller than those of their respective parent strains (Figure 6B and C, *p* < 1.28 x 10^-3^ and *p* < 3.06 x 10^-4^, respectively). *hv*Δr*osR* showed similar motility levels to that of the negative control strain, Δ*arlA[1-2]*, which was previously shown to be non-motile (Figure 6B). Taken together, these results suggest that RosR is necessary for motility in *Hfx*. species.

**Figure 6.**
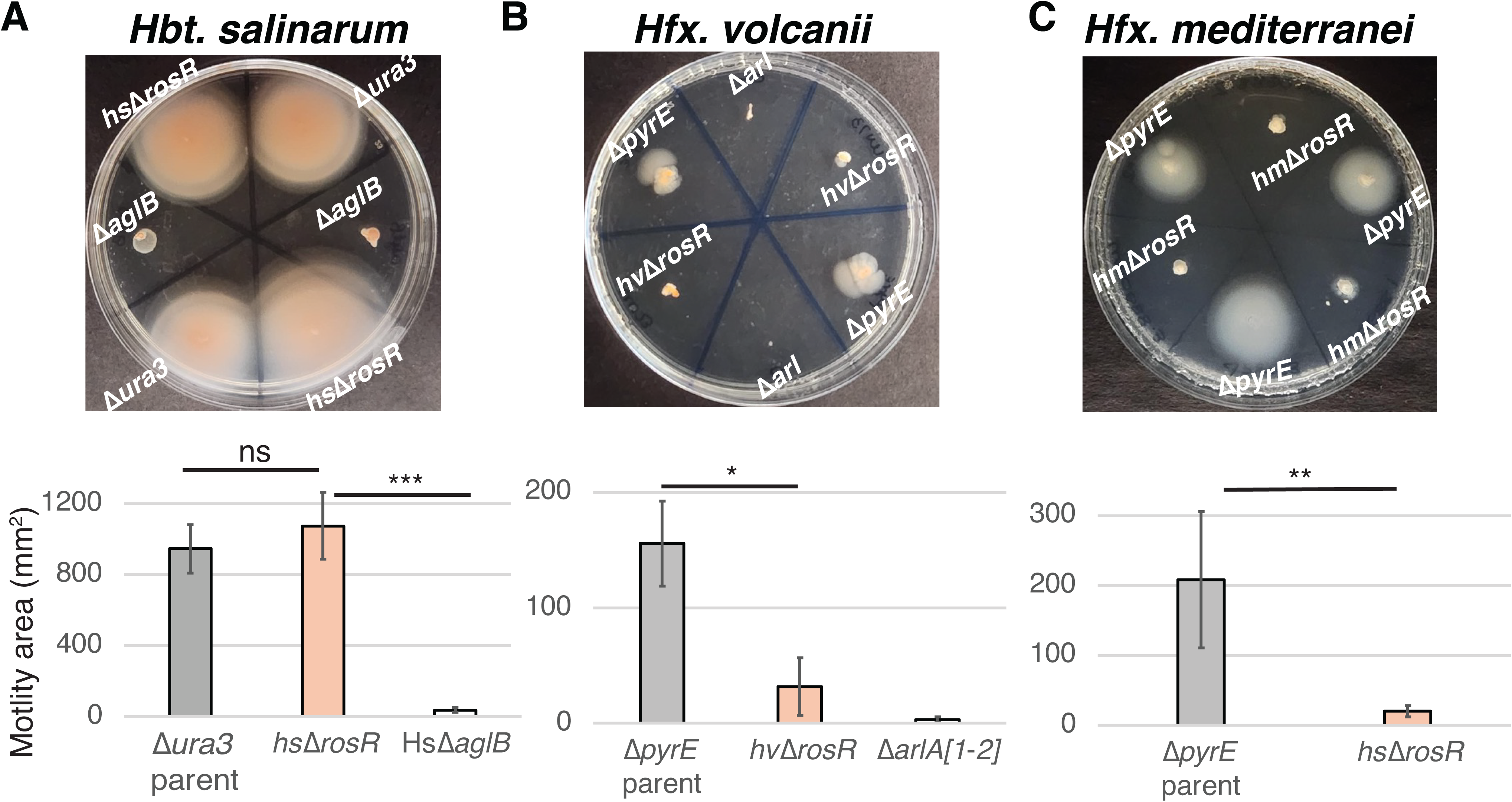
RosR is necessary for motility in *Hfx.* species. Original motility plates for a representative replicate are shown at top and bar graph quantification of motility halo area at bottom. Bar heights represent the average motility halo area (in mm^2^) of three replicate plates, each with two or three technical replicate spots. Error bars represent the standard deviation from the mean. Overbars indicate the statistical significance of the difference between the motility areas of each strain according to t-test: ns, not significant; ***, p < 10^-5^; **, p < 10^-4^; *, p < 10^-3^. See Supplementary Figure S2 for all biological replicate images.

### Flexibility of RosR-DNA binding may lead to rapid cis-trans rewiring

Given the strong amino acid sequence conservation in RosR protein regions that bind DNA (Figure 1B) but degeneracy in the cis-regulatory binding sequences across species (Figure 4, 5, (Kutnowski *et al*., 2019)), we reasoned that RosR from one species may be able to bind the cis-regulatory DNA motif of another. We therefore conducted AlphaFold3 modeling to predict and compare RosR-DNA structural interactions in all-by-all combinations across species. The AlphaFold model of the *hs*RosR DNA structure was strongly predictive of the known structure solved by X-ray crystallography (TM-score 0.95, RMSD 0.24, (Kutnowski *et al*., 2019)), suggesting that the model predicts the correct structure with high confidence (Figure S3A). Surprisingly, RosR from each *Hfx*. species was predicted to bind the known *hs*RosR binding sequence, yielding high confidence structural predictions (Figure S3B, top row, >90%). Similarly, high confidence predictions were observed for every other predicted RosR-DNA interaction, with slightly lower confidence for *hs*RosR interacting with the cis-regulatory sequences discovered in this study for *Hfx*. species (>80%, Figure S3, first column). In each case, RosR bent the DNA as expected from previous structural studies (Kutnowski *et al*., 2019). In each combination tested by AlphaFold, the wing region of RosR and the unstructured N-terminus of the protein gave lower confidence predictions, which may be indicative of higher protein flexibility. Although these models require future experimental validation, they suggest with high confidence that RosR of each species is capable of binding to the predicted cis-regulatory sequences of each other species. Together with the genomics data presented here (Figures 4 and 5), we interpret this result to mean that RosR may rewire through flexible cis-trans interactions.

## DISCUSSION

Here we report the surprising finding that RosR regulates motility and membrane genes in *Hfx. mediterranei* and *Hfx. volcanii.* This function stands in contrast to our previous findings that RosR in *Hbt. salinarum* functions as a global regulator of oxidative stress response (Sharma *et al*., 2012; Tonner *et al*., 2015). The RosR role in motility in the *Hfx.* species is based on the lack of growth defect in Δ*rosR* strains under oxidative stress conditions (Figure 2 and 3), the direct regulation of genes with motility and membrane-related functions (Figure 4 and 5), and the motility defect in *hm*Δ*rosR* and *hv*Δ*rosR* strains (Figure 6). This result is unanticipated given the strong primary sequence conservation of RosR throughout haloarchaea (Figure 1A), with extremely high conservation in the DNA binding helix and wing of the RosR protein across the species of interest here (Figure 1B).

Across *Hfx.* species, RosR plays a role in a complex network of regulators to govern archaeallum production and motility. In addition to the gene encoding ArlA2 in *Hfx. volcanii* (Figure 4), CirA, whose transcript is convergently transcribed with *arlA2,* was recently identified as another regulator of motility in *Hfx. volcanii* (Chatterjee *et al*., 2024). CirA exhibits 97% sequence identity to *Hfx. mediterranei* HFX_1222, with strong conservation of gene synteny in the motility gene cluster (Oberto, 2013). However, *hm*RosR was not observed to regulate HFX_1222 or the *arlA2* homolog HFX_1220 (Figure 5). Conversely, in *Hfx. mediterranei* HFX_1425, encoding a PGF domain containing protein, is regulated by *hm*RosR but its *Hfx. volcanii* homolog is not regulated by *hv*RosR. HFX_1425 encodes a putative outer membrane protein, an archeosortase substrate homologous to *Hfx. volcanii* HVO_2160 (rod determining factor A, RdfA). RdfA is a predicted substrate of the *Hfx. volcanii* ArtA archeosortase (Abdul Halim *et al*., 2013) and was recently shown to play a role in *Hfx. volcanii* cell shape regulation (Schiller *et al*., 2024). ArtA cleaves proteins containing a PGF motif for membrane lipid anchoring, including the cell surface layer (S-layer) glycoprotein (Abdul Halim *et al*., 2013; Pohlschroder *et al*., 2018). It is therefore likely that, by way of homology, HFX_1425 is also an ArtA substrate and outer membrane protein in *Hfx. mediterranei*. However, little is known regarding the mechanisms of motility in *Hfx. mediterranei*. In *Hfx. volcanii*, pilins may serve as posttranslational motility regulatory factors (Esquivel and Pohlschroder, 2014), and other putative motility regulatory mutants have also recently been identified (Collins *et al*., 2020). Together these results are consistent with the hypothesis that the concerted action of a complex network of factors, including RosR, regulate motility in the *Hfx*. species. It will be interesting in future research to compare the mechanisms of interplay between RosR, CinA, and other factors in the motility and cell surface regulatory network across species.

We hypothesize that DNA sequence discrimination by RosR is highly flexible across the three species of interest here. This flexibility may enable rapid alteration in target gene function over short evolutionary time scales. Previous structural studies suggested that *hs*RosR is flexible in its binding site sequence discrimination. Only two residues per monomer of the *hs*RosR dimer make hydrogen bonding contacts with DNA residues: H50 in the DNA binding helix and R74 in the wing region (Kutnowski *et al*., 2018; Kutnowski *et al*., 2019). These amino acid residues are completely conserved across RosR orthologs in the species studied here (Figure 1). Further, the *hs*RosR cis-regulatory DNA binding sequence was shown to be degenerate, with only three consensus residues per half site (TGT-N10-ACA)(Kutnowski *et al*., 2019). These repeated trimers are relatively weakly conserved in the predicted *Hfx. mediterranei hm*RosR binding sequence discovered here (Figure 5D), and only one of the two half-sites is conserved in *Hfx. volcanii* (Figure 4D). Therefore, even single base pair substitutions in non-coding DNA have the potential to completely change the location of RosR binding sites throughout the genome, thereby swapping RosR gene targets and altering the composition of the RosR regulon. Structural modeling showed that RosR from each of the three species studied here is predicted to be able to bind the putative cis-regulatory sequence from the other species, producing high confidence structural predictions (Figure S3). *hs*RosR was previously shown to bend DNA by 21.75° (Kutnowski *et al*., 2019), which was recapitulated by AlphaFold (Figure S3). How RosR discriminates between target sites to gain specificity remains to be determined, although it is possible that DNA structural features and/or other TF binding partners may play a role given RosR’s ability to bend DNA. Taking this evidence together, we hypothesize that RosR-DNA binding was rapidly rewired during evolution via flexibility in cis-trans interactions. This is supported by: (a) the predicted promiscuity of RosR binding to varying DNA cis-regulatory sequence across species (Figure S3); (b) the degeneracy in the cis-regulatory binding sequence, which could lead to a change in gene targets with a single SNP (Figure 5A and B, (Kutnowski *et al*., 2019)); and (c) the potential for RosR protein conformational flexibility (Figure S3, (Kutnowski *et al*., 2019)). The *Hfx*. species diverged ∼80 million years ago (Lopez-Garcia *et al*., 1995). Horizontal gene transfer is also common across *Hfx*. species, to the point that they can form stable interspecific hybrids (Naor *et al*., 2012). Together with the observed flexibility in RosR-DNA interactions and recent radiation of the haloarchaeal RosR protein clade (Figure 1A), RosR network rewiring may have outpaced the time of evolutionary divergence between the *Hfx*. genomes themselves, leading to a swap in RosR gene targets while maintaining the emergent property of motility regulation.

The three species, and haloarchaea more generally, dominate the biomass in evaporitic high desert hypersaline ecosystems such as salt flats and inland hypersaline lakes (Dead Sea, Great Salt Lake)(Schmid *et al*., 2020). However, *Hbt. salinarum* and the *Hfx.* species occupy different niches within these extremely saline systems (Andrei *et al*., 2012), which may have forced rapid RosR network rewiring. *Hfx*. species can be isolated from saline sediment and/or shore rock pools (Mullakhanbhai and Larsen, 1975; Turgeman-Grott *et al*., 2019). They grow optimally with fast doubling times in ∼2.5 M sodium chloride (Figures 2 and 3) and transition between biofilm and motile foraging lifestyles depending on oxygen availability and cell density (Hackley *et al*., 2024a; Schiller *et al*., 2024). In contrast, *Hbt*. *salinarum* has been detected primarily in hypersaline surface water, where it experiences high levels of UV and desiccation / rehydration cycles. *Hbt. salinarum* grows optimally with high sodium (∼4.3M NaCl) and relatively lower magnesium, which reduces the oxygen solubility. Its growth rate is slower than that of the *Hfx*. species (Figures 2 and 3), and less flexible in its lifestyle changes (constitutively rod-shaped and motile). In the case of RosR, differences in oxygen availability, reactive oxygen species (ROS) exposure, and differential ROS resistance (Figures 2 and 3) may play key roles in selection for rewiring. More generally, which specific selective force(s) drive TRN rewiring across species within these different selective regimes is an important avenue for future research.

In summary, we find that genes under RosR control change substantially over short evolutionary timescales in haloarchaea. Our data are consistent with the hypothesis that this rewiring occurs through flexibility in cis-trans interactions coupled with differential environmental selection. This finding is in contrast to other recently reported rewiring events in archaea such as the TrmB and DtxR transcription networks, which appeared to rewire via loss and gain of TFs while maintaining strong conservation of cis-regulatory binding sequences (Martinez-Pastor *et al*., 2017a; Hackley *et al*., 2024a; Hackley *et al*., 2024b; Martinez Pastor *et al*., 2024). Therefore, archaeal TRNs may be capable of rewiring using mechanisms similar to both those of eukaryotes (Tuch *et al*., 2008; Johnson, 2017) and bacteria (Perez and Groisman, 2009b, a).

## MATERIALS AND METHODS

### Media, routine growth conditions, and strain construction

Wild-type strains *Halobacterium salinarum* NRC-1 (*Hs)*, *Haloferax volcanii* strain DS2 (*Hv)*, and *Haloferax mediterranei* (*Hm)* ATCC 33500 were used in this study. The *Escherichia coli* cloning strain NEB5α (New England Biolabs) was used for propagation of plasmids. *Hs* strains were grown in complete medium (CM) which contained 250 g/L NaCl (Fisher Scientific), 20 g/L MgSO_4_·7H_2_O (Fisher Scientific), 3 g/L trisodium citrate (Fisher Scientific), 2 g/L KCl (Fisher Scientific), 10 g/L Bacteriological Peptone (Oxoid), pH 6.8. *Hm* and *Hv* strains were grown in Hv-YPC medium, which contained per liter: 144 g NaCl, 21 g MgSO_4_·7H_2_O, 18 g MgCl_2_·6H_2_O, 4.2 g KCl, and 12 mM Tris HCl (pH 7.5). Hv-YPC medium was prepared as instructed in (Allers *et al*., 2004). All media were supplemented with uracil to complement the biosynthetic auxotrophy of the parent strain (50 µg/ml) and species routinely grown at 42°C.

The gene encoding RosR was deleted using suicide integrative plasmids specific for each species (HVO_0730 in *Hv*, and HFX_0688 in *Hm*). The *Hs* VNG_0258H deletion strain (*hs*Δ*rosR*) was constructed previously (Sharma *et al*., 2012). In each of the *Hfx.* species, strains encoding RosR tagged with the hemagglutinin (HA) epitope at the C-terminus were constructed by integrating the tag into the genome at the native locus. The *Hbt.* RosR epitope-tagged strain was constructed previously (Tonner *et al*., 2015). To generate these strains, deletion and integrant plasmids were constructed using isothermal ligation (Gibson *et al*., 2009). Specifically, PCR products were generated using Phusion DNA polymerase (New England Biolabs) and assembled to plasmid backbones linearized with restriction enzyme(s) (Table S2). In general, plasmids were transformed into each of the three species using established protocols (Peck *et al*., 2000; Allers *et al*., 2010; Liu *et al*., 2011), except that halophile strains were transformed directly with NEB5α produced plasmids. Double crossover counterselection methods were used to generate deletion and HA-tagged strains. For selection of first crossover events during strain construction (“pop-in”), *Hs* was selected on CM agar containing uracil (50 µg/ml) and mevinolin (10 µg/ml). *Haloferax* strains were selected on HvCA agar (Allers *et al*., 2010). Counterselection (“pop-out”) for deletion strains was performed in the presence of uracil and 5-Fluoroorotic acid (*Hs* 300 µg/ml, *Hm*: 250 µg/ml, *Hv* 50 µg/ml). All strains were verified by PCR and Sanger sequencing at the junction. Primers were obtained from Integrated DNA Technologies (Coralville, IA). The Δ*rosR* strains and *Hfx. mediterranei* WR510 (Δ*pyrE2* parent) were validated by whole genome resequencing as described in (Zaretsky *et al*., 2019) and in the Github repository https://github.com/CindyDarnell/aglB-WGS-growth. The whole genome of *Hfx. volcanii* Δ*pyrE* parent strain (H26) used here was previously validated (Soborowski *et al*., 2024). Raw sequence reads are freely available for download at the NCBI Sequence Read Archive (SRA) via accession number PRJNA1173512. All strains, primers, and plasmids are listed in Supplementary Table 2.

### Sequence and structural bioinformatic analyses

All protein sequences for PF03551 (PadR-like family) were downloaded from pfam.xfam.org (doi: 10.1093/nar/gky995). Taxonomy and gene locus information was downloaded from uniport.org (doi: 10.1093/nar/gky1049). Data were downloaded in May 2019. After removing deprecated sequences, the final sequence file contained 767 protein sequences (Supplementary Table 1). Sequences were aligned using clustal Omega with default settings for neighbor joining method using the Gonnet protein weight matrix (Sievers *et al*., 2011; Sievers and Higgins, 2018). The resulting tree was visualized using the interactive Tree of Life website (itol.embl.de; (Letunic and Bork, 2019)). The RosR clade was identified by assessing synteny with the *ipp*/*ppa* gene using the NCBI genome browser for each locus from a haloarchaeal genome. Percent identity for each residue in the PadR domain of the 91 sequences in the RosR clade was calculated against the consensus sequence after alignment. A protein weblogo was created from the same alignment (Crooks *et al*., 2004). For protein-DNA structural predictions, the AlphaFold3 server (alphafoldserver.com) was used with default parameters (Abramson *et al*., 2024). All-by-all combinations of *hs*RosR, *hv*RosR, or *hm*RosR as protein input was compared to cis-regulatory binding sequence for each species. The TM-align algorithm (Zhang and Skolnick, 2005) available at the Protein Data Bank RCSB Pairwise structural alignment tool (https://www.rcsb.org/docs/tools/pairwise-structure-alignment) was used with default parameters compare the published *hs*RosR-DNA structure (PDB 6QIL) to the AlphaFold3 predicted structure.

### High throughput growth curve experiments and analysis

Strains were grown to early stationary phase in 3 ml media and subcultured to an optical density at 600 nm of 0.05. Multi-well plate growth curves were performed in a Bioscreen C microbial growth analyzer in 100-well Honeycomb plates (Growth Curves USA, Piscataway, NJ) set to measure OD600 every 30 minutes. Plates were incubated at 42°C. H_2_O_2_ (Sigma) and paraquat (methyl viologen dichloride) (Sigma) and were added at time zero at concentrations indicated. Growth curve data (Table S3) was plotted using the tidyverse package in R in the RStudio framework. Code and package version information are provided freely on the github repository associated with this study (https://github.com/amyschmid/rosr_3species). Area under the curve (AUC) metric was calculated as previously described in (Todor *et al*., 2014), with code available at https://github.com/wleepang/BioscreenUtils. Significant differences in AUC between strains were calculated using the Welch’s two-sided t-test in R. AUC plots for each species, strain, and condition are given in Figure S1C.

### ChIP-seq and ChIP-qPCR experimental and analytical methods

Each *Hfx.* strain (parental Δ*pyrE* and RosR::HA tagged) was grown in triplicate (ChIP-qPCR) or quadruplicate (ChIP-seq) in 50 mL rich medium until mid-to late-exponential phase (OD_600_ ≍0.5-1.0). Crosslinking was performed by adding 37% formaldehyde (CH_2_O) for 20 min at room temperature and chromatin immunoprecipitation (ChIP) was conducted as previously described (Martinez-Pastor *et al*., 2017a).

For ChIP-seq, libraries were constructed using the KAPA Hyper Prep kit using Illumina TruSeq adapters. DNA library quality was assessed by Bioanalyzer using a High Sensitivity DNA chip (Agilent). Samples for each species were multiplexed according to the adapter mixtures and pooled. Each pool was run in a single lane on an Illumina HiSeq 4000 instrument by the Duke Sequencing and Genomics Technologies core, yielding single-end sequence reads. 50 bp reads were assessed for quality using FastQC and adapter sequences trimmed using Trim Galore and Cutadapt. Resultant reads were mapped to the *Hfx. mediterranei* ATCC3500 (assembly ASM30676v2; NCBI accession GCF_000306765.2) or *Hfx. volcanii* DS2 genome (ASM2568v1; GCF_000025685.1) using Bowtie2. Sorting of mapped sequences was conducted using samtools. Sites of significant RosR binding enrichment were called using the MOSAiCS peak-calling package (Sun *et al*., 2013) with sorted bam files as input. MOSAiCS parameters included fragment length (fragLen) 200, cap 0, PET false, analysis type IO, background estimate rMOM, FDR 0.01. Using the DiffBind R package (Ross-Innes *et al*., 2012), peaks were considered reproducible if 100 bp of the reads in the peak overlapped across three of four biological replicate experiments. DNA within peak regions was used as input to the MEME suite for motif detection, then whole genomes of each species were scanned for additional motifs using FIMO within the MEME suite (Bailey *et al*., 2015). Only motif instances with FIMO q-value < 0.85 were accepted as significant (Table S4). Using the GenomicRanges and IRanges R packages (Lawrence *et al*., 2013), a given ChIP-seq peak was associated with a given genomic feature (gene or promoter region) if they overlapped by 100 bp. Peaks that fell entirely inside promoter regions were associated with the nearest gene within 500 bp. Peak-to-gene associations were then annotated using the corresponding NCBI genome annotation within the GenomicFeatures R package. Peaks were plotted and further validated visually using the trackvieweR package and the Integrated Genome Viewer (Robinson *et al*., 2023). trackvieweR visualizations are shown in the Figures. ChIP-seq raw and processed data are freely available for download at the National Center for Biotechnology Information (NCBI) Gene Expression Omnibus (GEO) accession GSE279075. Code and package version information are provided freely on the github repository associated with this study (https://github.com/amyschmid/rosr_3species).

For ChIP coupled to quantitative PCR (ChIP-qPCR), DNA purified from ChIP reactions was analyzed in technical triplicate by real time qPCR (Roche LightCycler^®^ 96 System) using dye-based qPCR master mix according to manufacturer’s instructions (VWR^R^ qPCR Master Mix). Three primer sets (Integrated DNA Technologies, Coralville, IA, USA) were designed to amplify the region enriched for RosR binding in ChIP-seq experiments, namely, in the region between genes HVO_1211 (*arlA2*) and HVO_1212 (*cirA*). A control primer pair was designed in the region just upstream of HVO_1211. Primer sequences are detailed in Supplementary Table 2 and a schematic representation of the position of the primers is depicted in Figure 4. Primer efficiency was calculated using serial dilutions of genomic DNA before the enrichment analysis. DNA binding enrichment in the region of interest relative to the control region was calculated from primer efficiency and threshold crossing point (Ct) values of each of the three amplicons (peak1, 2 and 3) compared to the control amplicon using the Pfaffl method (Pfaffl, 2001; Martinez-Pastor *et al*., 2017a). Values of three biological replicates and three technical replicates were averaged and standard error of the mean calculated across biological replicates. Statistical significance reported in the text and figure was determined by two-sided unpaired Student’s t-test.

### RNA-seq experimental and analytical methods

*hm*Δ*rosR*, *hv*Δ*rosR*, and corresponding parent strain cultures were grown in biological triplicate in rich medium to mid-exponential phase. RNA was extracted using an Absolutely RNA Miniprep kit (Agilent, Santa Clara, CA). RNA quality was measured using RNA 6000 Nano kit on a 2100 Bioanalyzer (Aglient, Santa Clara, CA). PCR to verify lack of DNA contamination was conducted on 300 ng of RNA sample using species-specific primers listed in Supplementary Table S2. Ribosomal RNA depletion was performed from 300 ng of each sample using NEBNext^R^ RNA Depletion Core Reagent (#E7870 New England Biolabs) according to the manufacturer instructions using probes reported in (Pastor *et al*., 2022). The resultant mRNA was purified using NEBNext RNA sample purification beads and eluted in 7 μl of nuclease-free water. 1μl of each sample was used for quantification and quality verification (free of rRNA) in a second round of Agilent Bioanalyzer analysis using 2100 Prokaryote Total RNA Nano kit (Agilent, Santa Clara, CA). Libraries were constructed using 5 μl of mRNA using the Ultra II Directional RNA Library Preparation Kit (Agilent, Santa Clara. CA). Indexes were designed according to the kit specifications and obtained from Integrated DNA Technologies. Resultant libraries were tested for quality and quantity by Bioanalyzer using the 2100 expert High Sensitivity DNA Assay chip (Agilent Technologies) and sequenced on an Agilent NovaSeq6000 instrument and data analyzed by the Duke Genomics and Sequencing Technologies core facility.

RNA-seq data were processed using the fastp toolkit (Chen *et al*., 2018) to trim low-quality bases and sequencing adapters from the 3’ end of reads. STAR (Dobin *et al*., 2013) RNA-seq alignment tool was used to map trimmed reads to the genome of *Haloferax mediterranii* or *Haloferax volcanii* (using NCBI assemblies ASM30676v2 or ASM2568v1, respectively). Reads aligning to a single genomic location were summarized across genes. For genes having an overlap of at least 10 reads, gene counts were normalized and differential expression was carried out using the DESeq2 (Love *et al*., 2014) Bioconductor (Huber *et al*., 2015) package implemented for the R programming environment. Consistent with the recommendation of the DESeq authors, independent filtering (Ignatiadis *et al*., 2016) was utilized prior to calculating adjusted p-values (Benjamini Y, 1995) and moderated log2 fold-changes were derived using the ashr package (Stephens, 2017). Genes were considered significantly differentially expressed if the log2 fold change ratio (Δ*rosR* : parent) was >= |1| and adjusted p-value < 0.05. RNA-seq raw and processed data are freely available for download at NCBI GEO via accession GSE279016. Code and package version information are provided freely on the github repository associated with this study (https://github.com/amyschmid/rosr_3species).

### Motility assays

The parental strains, Δ*rosR* mutants, and known non-motile control strains (MT2, AKS211, Table S2) from the three species studied in this work were streaked from -80°C freezer stocks onto rich media plates and incubated for 5-10 days at 42°C until single colonies were visible. Two to three colonies from each strain were then stabbed into HvCa18%ura^+^ (for *Hfx.* strains) or CMura^+^ (*Hbt. salinarum*) soft agar (0.35%) plates. The media recipes for these plates are described above. The plates were incubated statically at 42°C for 4 days (*Hfx. mediterranei*), 6 days (*Hfx. volcanii*), or 10 days (*Hbt. salinarum*) before imaging. Images were captured using a S22+ camera (10MP F2.2 [Dual Pixel AF], FOV 80°, 1/3.24”, 1.22µm) within an imaging box constructed from cardboard and black paper, following recommendations from (Smith and Schuster, 2021). To calculate colony area, we utilized ImageJ free software (Schneider *et al*., 2012), setting the scale for measurements based on the known plate diameter (88.9 mm). Images were standardized to 16-bit and subjected to fixed color thresholding. Colony area was determined using the “measure” command in the ImageJ program graphic user interface “Analysis” menu (Ducret *et al*., 2016). Data from two or three colonies per plate across three different plates were averaged and represented in Excel. Statistical significance (*) was assessed using a two-tailed, two-sample unequal variance t-test.

## ACKNOWLEDGEMENTS

The authors thank Schmid lab members past and present (especially Rylee Hackley, Sungmin Hwang, Alex Phillips) for comments on the study during its development. Thank you to Alexis Bailey and Andrew Soborowski for comments on the manuscript. We thank Joey Prinz and Devi Swain-Lenz in the Duke Sequencing and Genomics Technologies core facility for technical support with sequencing and RNA-seq data analysis. This study was funded by National Science Foundation grants 1651117, 1936024, and 2427099 to AKS.

## GUIDE TO SUPPLEMENTARY MATERIAL

**Supplementary Figure S1.** Raw growth data plots. Each point represents one time point for one culture (well) of the high throughput plate reader. (A) H_2_O_2_ and PQ titration data as shown in main text Figure 2. (B) Comparison of growth phenotypes between parent strain and Δ*rosR* mutants across species. Colors and oxidant concentrations correspond to main text Figure 3. (C) Comparison of average area under the curve (AUC) for each species, strain, and condition. Bars represent the average AUC for at least 12 replicate trials, error bars the standard deviation from that mean. “NS” overbar for *Hfx. volcanii* under 0.5 mM H_2_O_2_ depicts that the parent strain AUC is not significantly different than the Δ*rosR* mutant strain by Welch’s two-sample t-test.

**Supplementary Figure S2.** Original images of all replicates of motility plates and corresponding thresholded images processed in ImageJ (red color does not have a meaning). Species are given in the columns and biological replicates in the rows. Plate position labels for each strain are given at top and are consistent throughout all the images for a given species.

**Supplementary Figure S3.** AlphaFold3 (Abramson *et al*., 2024) structural model predictions of RosR-DNA interactions. Table shows all-by-all combinations of RosR-DNA sequence binding from each species. *Hbt. salinarum* (*hs*RosR), *Hfx. volcanii* (*hv*RosR) and *Hfx. mediterranei* (*hv*RosR) are shown in the columns, whereas each predicted cis-regulatory DNA binding site is in the rows. Cis-regulatory sequence logos shown for Hfx. volcanii and Hfx. mediterranei are also shown in Figures 4 and 5, respectively, and discussed in the main text. Specifically, DNA sequences used for structural modeling are: HbtSELEX, GCGAGG**TGT**AAATTGTCTG**ACA**TGTTCT, bold residues correspond to those that were structurally determined to contact RosR residues (Kutnowski *et al*., 2019); Hvo predicted, GCGTAA**TCA**AATTTCTG**ACA**TTAA, predicted by MEME (see Methods); Hme predicted, ATTAAT**TGA**CAATTTATTG**ACA**TGTTAA. Dark blue shading of protein-DNA structures in Fig S3 indicates high confidence predictions, whereas residues trending from cool colors toward hot colors indicates decreasing confidence (see color scale bar at bottom right). Binding predictions were made using the freely available AlphaFold3 Google server at https://alphafoldserver.com/ using default parameters, accessed on August 9, 2024.

