## Supplementary figures and images for "Rapid rewiring of an archaeal transcription factor function via flexible cis-trans interactions"

### Supplementary Figure 1

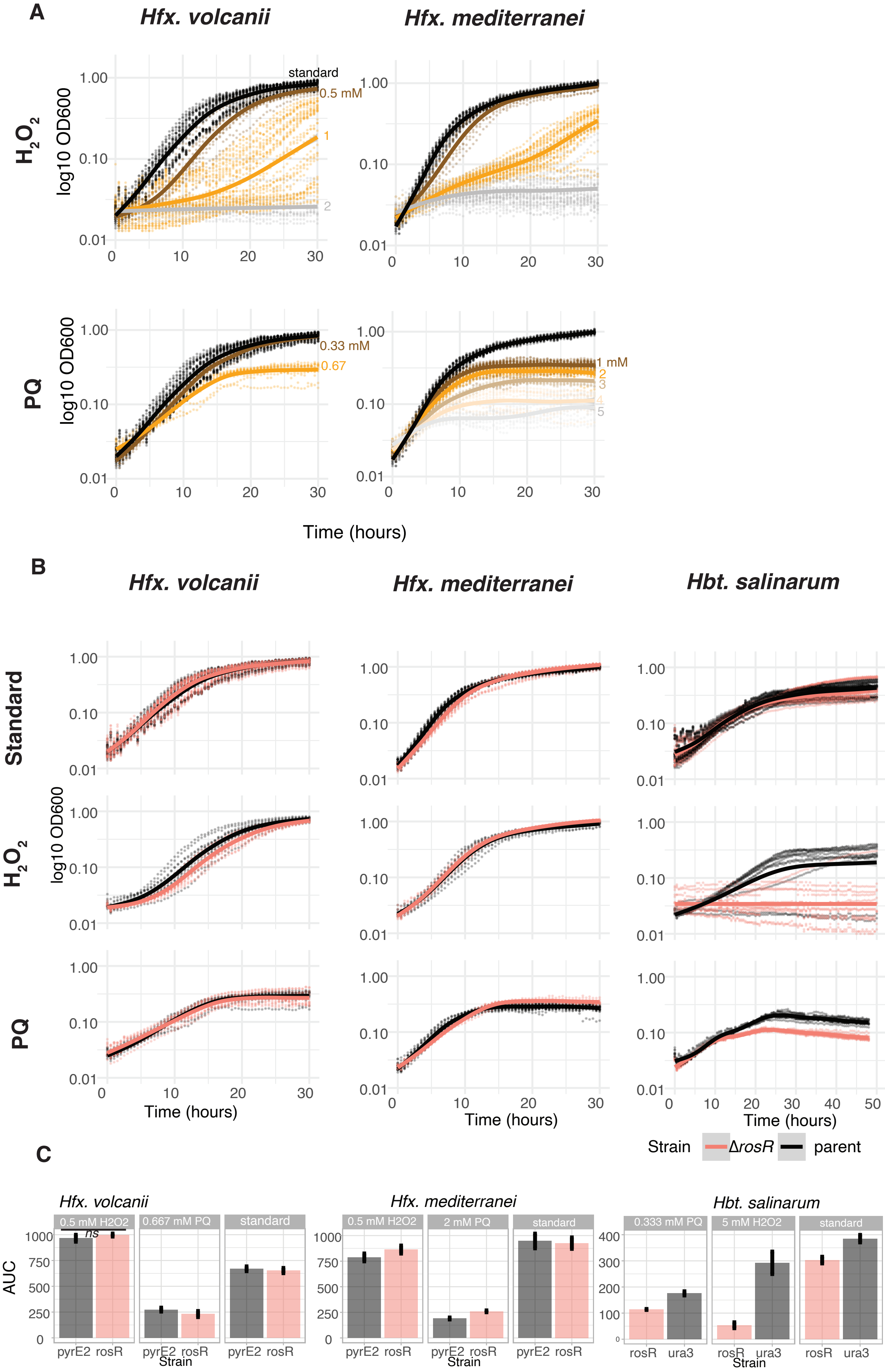

### Supplementary Figure 2

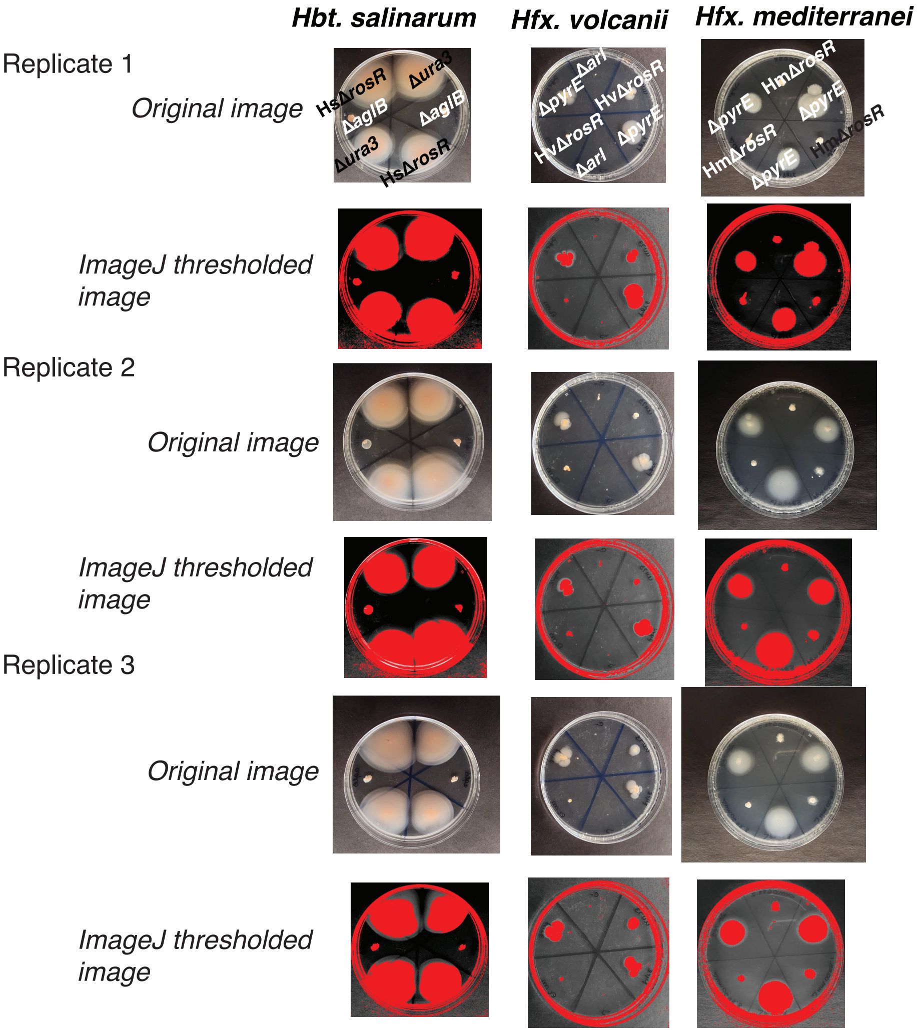

### Supplementary Figure 3

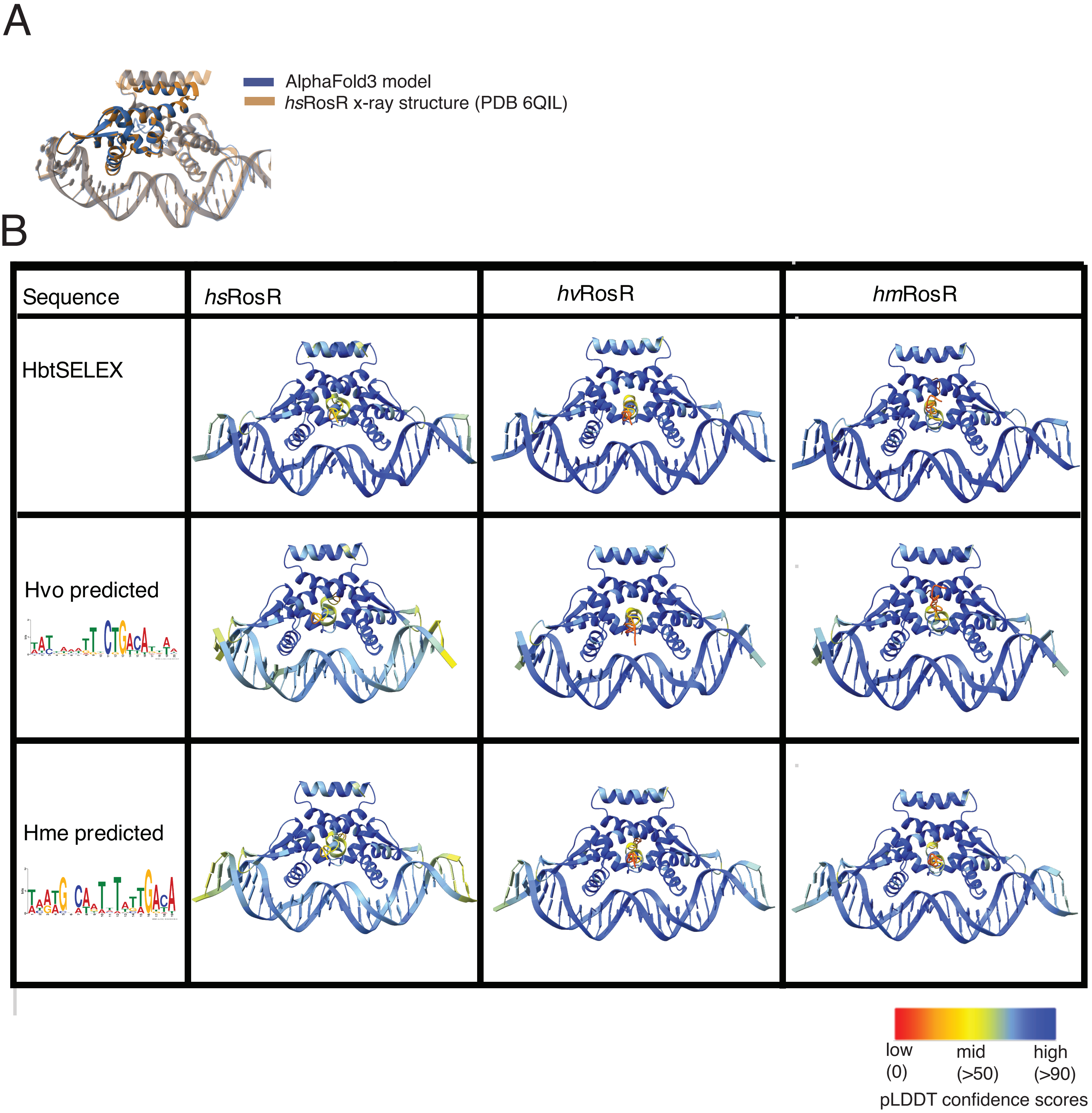
